# Practical and Thermodynamic Constraints on Electromicrobially-Accelerated CO_2_ Mineralization

**DOI:** 10.1101/2022.01.09.475503

**Authors:** Sabrina Marecos, Rae Brigham, Anastacia Dressel, Larissa Gaul, Linda Li, Krishnathreya Satish, Indira Tjokorda, Jian Zheng, Alexa M. Schmitz, Buz Barstow

## Abstract

By the end of the century tens of gigatonnes of CO_2_ will need to be removed from the atmosphere every year to maintain global temperatures. Natural weathering of ultramafic rocks and subsequent mineralization reactions can convert atmospheric CO_2_ into ultra-stable carbonates. But, while natural weathering will eventually draw down all excess CO_2_, this process will need hundreds of thousands of years to do it. The CO_2_ mineralization process could be accelerated by weathering ultramafic rocks with biodegradable lixiviants like organic acids. But, in this article we show that if these lixiviants are produced from cellulosic biomass, the demand created by CO_2_ mineralization could monopolize the world’s supply of biomass even if CO_2_ mineralization performance is high. In this article we demonstrate that electromicrobial production technologies (EMP) that combine renewable electricity and microbial metabolism could produce lixiviants for as little as $200 to $400 per tonne at solar electricity prices achievable within the decade. Furthermore, this allows the lixiviants needed to sequester a tonne of CO2 to be produced for less than $100, even with modest CO_2_ mineralization performance.

## Introduction

The IPCC’s (Intergovernmental Panel on Climate Change) 2018 special report on the impact of climate change highlighted the need for significant deployment of negative emissions technologies to limit global warming [ipcc2018a]. The IPCC estimates that by the end of the 21st century, ≈ 20 gigatonnes of CO_2_ (GtCO_2_) will need to be removed from the atmosphere every year to limit global temperature rise to 1.5°C [ipcc2018a].

Of all the negative emissions technologies examined for large scale CO_2_ removal, carbon mineralization has the largest potential storage capacity [Keleman2019a, Beerling2020a, Lehmann2020a]. The CO_2_ storage capacity of carbon mineralization in ultramafic systems is truly enormous. For example, peridotite reservoirs across the globe (largely containing the mineral olivine) have the potential to mineralize and sequester 10^5^-10^8^ GtCO_2_ [Keleman2019a], between 100 and 100,000 × the excess CO_2_ in the atmosphere (there are ≈ 600 Gt more CO_2_ in the atmosphere than in pre-industrial times, and ≈ 430 Gt more CO_2_ in the oceans [NOAA2021a]).

Natural weathering of exposed sections of mantle rocks will eventually draw down all excess CO_2_ in the atmosphere, but will take thousands of years to do it [Archer2009a].

Mineral-dissolving microbes could accelerate mineral weathering. However, almost all mineral-dissolving microbes need to be powered by the degradation of plant biomass (*i*.*e*., the product of photosynthesis). For example, the mineral-dissolving microbe *Gluconobacter oxydans* B58 oxidizes the sugar glucose to the environmentally benign lixiviant (a mineral-dissolving compound) gluconic acid (glucose can be derived from degradation of cellulose, one of the primary components of biomass) [Reed2016a, Schmitz2021b].

However, the world’s growing [Prosekov2018a] and increasingly wealthy population [PwC2015a] is creating a growing need for arable land [Tilman2011a], tightening the world’s biomass supply [Slade2014a]. Could the use of plant biomass to power CO_2_ mineralization compete with the world’s food supply?

Electromicrobial production (EMP) could enable production of lixiviants for CO_2_ mineralization without competing with the world’s biomass supply. EMP technologies [Rabaey2010a, Rabaey2011a, Lips2018a, Salimijazi2019a, Claassens2019a, Prevoteau2020a] that combine biological and electronic components have been demonstrated at lab scale to have energy to chemical conversion efficiencies exceeding all forms of terrestrial photosynthesis [Liu2016a, Haas2018a], while theoretical predictions indicate that their efficiency could exceed all forms of photosynthesis [Claassens2019a, Salimijazi2020b, Leger2021a, Wise2021a]. Globally, photosynthesis has an average solar to biomass conversion of less than 1% [Barstow2015a]. In contrast, lab scale experiments have demonstrated a solar to product conversion efficiency of ≈ 10% for EMP [Liu2016a], while theoretical predictions indicate that this could rise to over 15% [Salimijazi2020b]. This order of magnitude increase in solar to product conversion efficiency could allow the production of lixiviants with no competition for arable land or wilderness.

In this article, we present a simplified model that estimates the global need for lixiviants for CO_2_ mineralization, the costs of synthesizing these lixiviants by electromicrobial production, and the costs of sequestering 1 tonne of CO_2_ using electromicrobially produced lixiviants.

## Theory

A full set of symbols used in this article is included in **Table 1**.

**Table 1.**
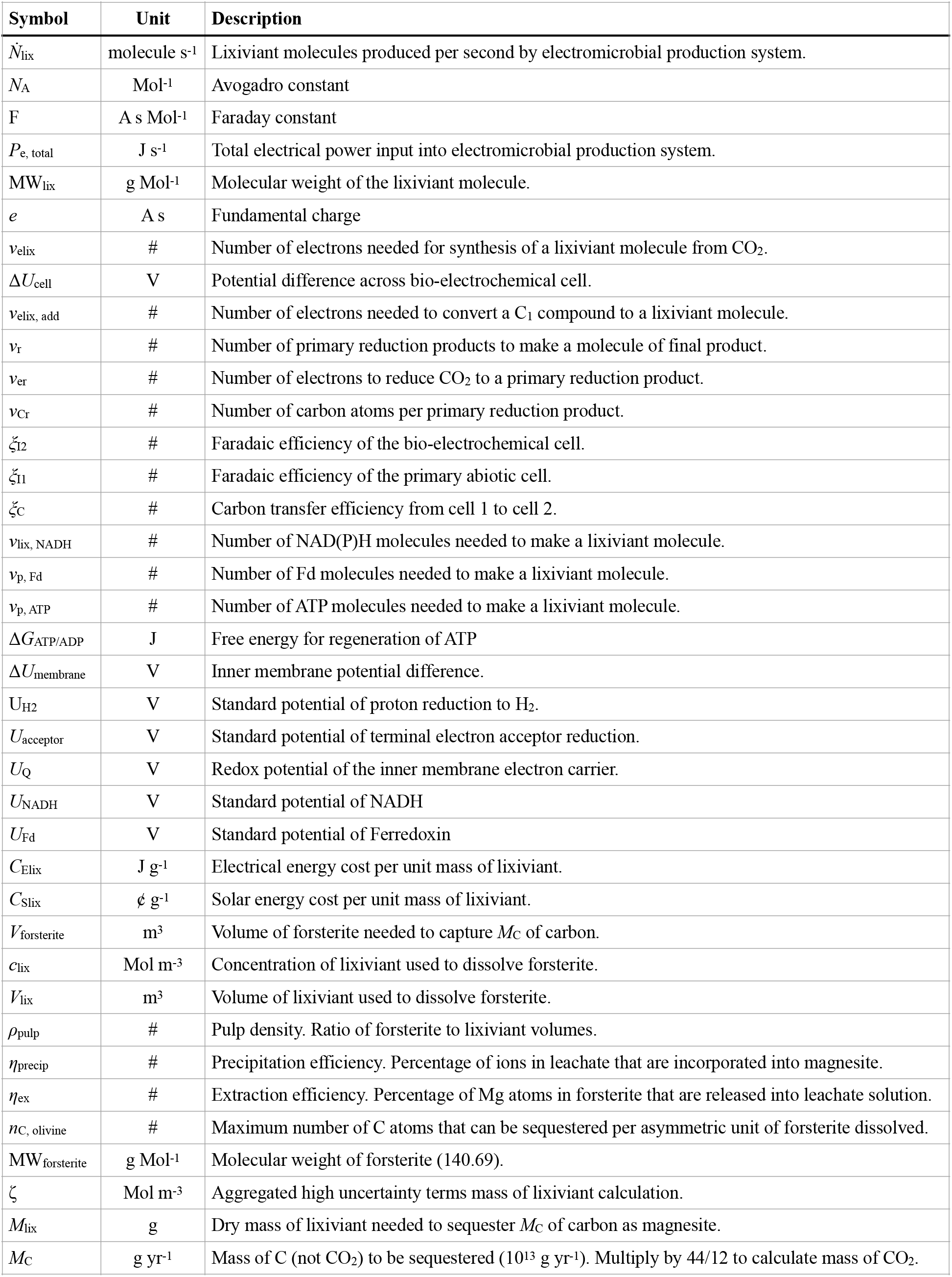
Symbols used in this article.

### Simplified Carbon Mineralization Reactions and Lixiviant Need

How much lixiviant is required to capture 20 GtCO_2_ per year (the approximate quantity estimated by the IPCC in order to limit global temperature rise to ≈ 1.5 °C [ipcc2018a])? To simplify the calculation, we consider just the conversion of magnesium olivine (forsterite) into magnesium carbonate (magnesite) through a two-step reaction. In the first step, solid forsterite is dissolved to aqueous (aq) magnesium ions [Power2013a],

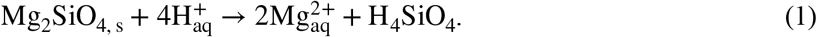

In a later precipitation reaction, these Mg^2+^ ions react with atmospheric CO_2_ and precipitate as stable solid (s) carbonates including magnesite (MgCO_3_) [Power2013a],

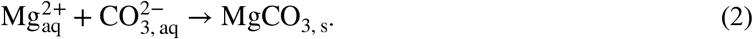

How much forsterite needs to be dissolved to capture 20 GtCO_2_? The maximum number of CO_2_ molecules (or C atoms) that can be sequestered by the dissolution of a single asymmetric unit of forsterite (Mg_2_SiO_4_), *n*_C, olivine_, is 2 (one asymmetric unit of forsterite contains 2 Mg atoms, which can each react with 1 carbon atom). The molecular weight of a single forsterite asymmetric unit is 141 grams per mole, and the molecular weight of 2 C atoms is 24 grams per mole. Thus, the minimum mass of forsterite needed to capture a mass of carbon *M*_C_ (*e*.*g*., 0.27 GtC corresponding to 1 GtCO_2_), is,

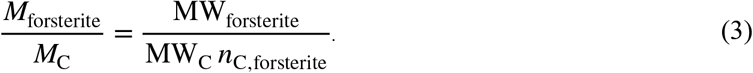

Therefore, to sequester 1 gigatonne of CO_2_, at least 16 gigatonnes of forsterite need to be dissolved [Power2013a].

How much lixiviant is needed to dissolve this much forsterite? The volume of the forsterite can be simply calculated from its density, *ρ*_forsterite_,

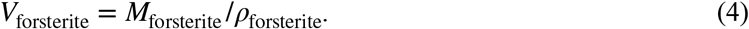

The volume of the lixiviant, *V*_lix_, can be calculated from the experimentally-derived pulp density that gives the best mineral dissolution,

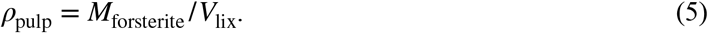

*ρ*_pulp_ is typically expressed in % w/v. For example, *ρ*_pulp_ = 2 %, means that 2 grams of forsterite are dissolved in 100 mL of lixiviant. However, so that we can use the experimentally derived pulp density along with our preferred units, we express *ρ*_pulp_ in terms of g m^−3^ (simply multiply *ρ*_pulp_ in w/v by 10^4^).

The mass of the dry lixiviant can be calculated simply from its molecular weight; concentration, *c*_lix_; and volume, *V*_lix_,

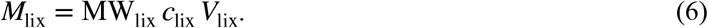

A full listing of molecular weights of the lixiviant compounds considered in this article is included in **Table S1**.

Thus, the minimum mass of the lixiviant needed to dissolve *M*_forsterite_, and hence to sequester *M*_C_ of carbon is,

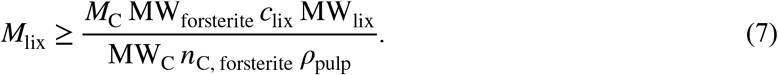

However, not all steps in the CO_2_ mineralization process will be perfectly efficient. The extraction of Mg from forsterite will be imperfect (**Equation 1**), as will the later precipitation of Mg^2+^ ions as a carbonate (**Equation 2**). To account for this, we introduce extraction efficiency, *η*_ex_, and precipitation efficiency, *η*_Precip_,

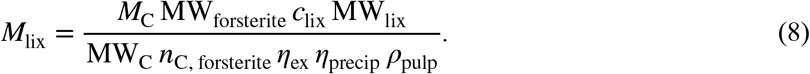

The formula for the mass of lixiviant, M_lix_, required to sequester a given amount of carbon per year, is composed of two sets of terms: those with at least reasonably well known values (MW_forsterite_, MW_C_, *n*_C, forsterite_), and a second set whose values have high uncertainty (*η*_ex_, *η*_precip_, *ρ*_pulp_, *c*_lix_),

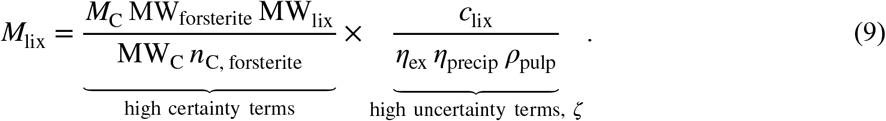

We denote the product of the high uncertainty terms as *ζ*, the inverse CO_2_ mineralization performance. The higher *ζ* gets, the more lixiviant it takes to sequester *M*_C_. Given that the uncertainty in each of the four terms in *ζ* is equally high, we choose to make our estimate of *M*_lix_ a function of *ζ* rather than any single uncertain parameter. Thus,

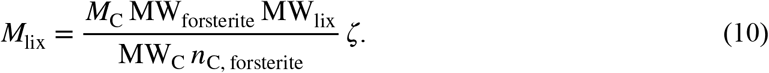

### Theory of Electromicrobial Production

We have extended our theoretical framework for calculating the efficiency of electromicrobial production (EMP) [Salimijazi2020b, Wise2021a] to calculate the energy cost of lixiviant production from renewable electricity and CO_2_. Full derivations of the equations presented here can be found in the supplement to our original electromicrobial production efficiency theory article (Salimijazi *et al*. [Salimijazi2020b]), and in our recent work on electromicrobial production of protein with extends our theory to calculate the energy (electrical or solar) costs of producing a gram of product (Wise *et al*. [Wise2021a]).

We consider a bio-electrochemical system used to deliver electrons to microbial metabolism (**Fig. 1B**). Electrical power is used to generate lixiviant molecules with a molecular weight MW_lix_. The amount of electricity needed to produce a unit-mass of the lixiviant is,

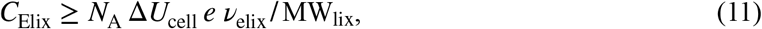

where *eν*_elix_ is the amount of charge needed to synthesize a single lixiviant molecule from CO_2_ (the fundamental charge, *e*, multiplied by the number of electrons needed for synthesis, ν_elix_); ΔU_cell_ is the potential difference across the bio-electrochemical cell; and *N*A is the Avogadro constant. A derivation of **Equation 11** can be found in Wise *et al*. [Wise2021a], building upon derivations in Salimijazi *et al*. [Salimijazi2020b].

**Figure 1.**
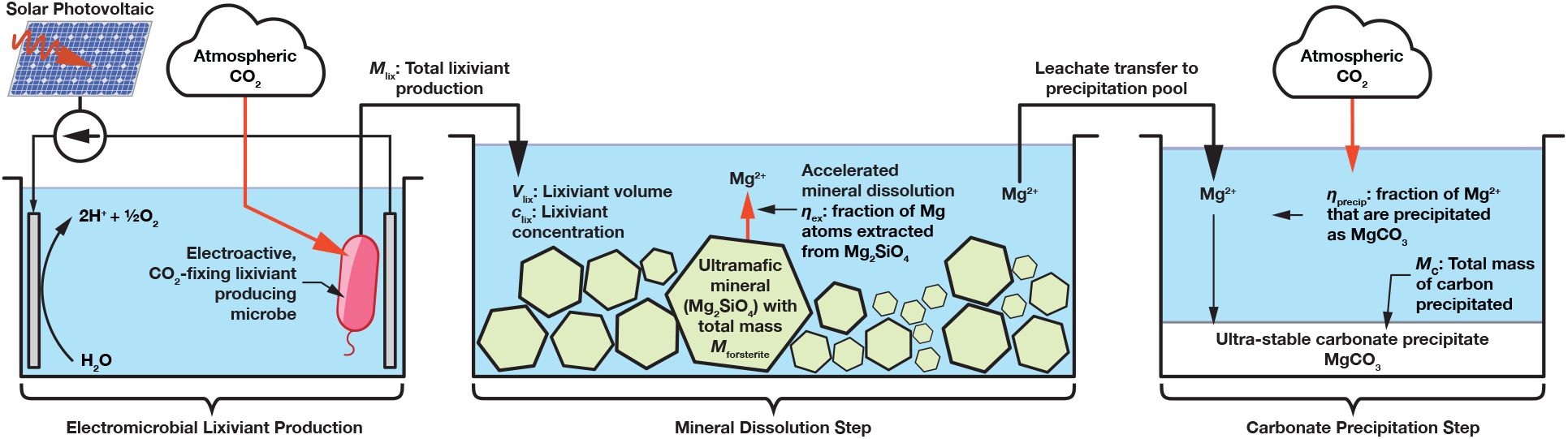
Overview of electromicrobially accelerated CO_2_ mineralization process. Key parameters in this article are highlighted in this figure, **Figure 2**, and **Tables 1** and **2**.

For systems where CO_2_ reduction is performed electrochemically, and the resulting reduction product (typically a C_1_ compound like formic acid) [Appel2013a, White2014a, White2015a] is further reduced enzymatically, *ν*_elix_ is substituted for number of electrons needed to convert the C_1_ product into the lixiviant, *ν*_elix, add_ [Salimijazi2020b],

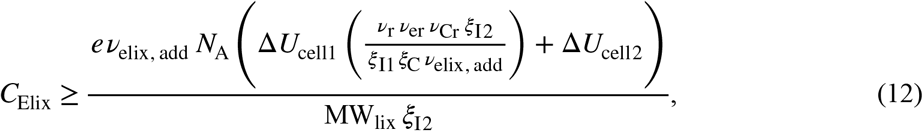

where *ν*_r_ is the number of primary reduction products (*i*.*e*., formic acid molecules) needed to synthesize a molecule of the final product, *ν*_er_ is the number of electrons needed to reduce CO_2_ to a primary reduction product (*i*.*e*., 2 in the case of formic acid), *ν*_Cr_ is the number of carbon atoms per primary fixation product (*i*.*e*., 1 in the case of formic acid), *ξ*_I2_ is the Faradaic efficiency of the bio-electrochemical cell, *ξ*_I1_ is the Faradaic efficiency of the primary abiotic cell 1, *ξ*_C_ is the carbon transfer efficiency from cell 1 to cell 2. A derivation of **Equation 12** can be found in Wise *et al*. [Wise2021a].

We calculate the electron requirements for lixiviant synthesis, *ν*_elix_ (from CO_2_) or *ν*_elix, add_ (from an electrochemical CO_2_ reduction product), from the number of NAD(P)H (*ν*_lix, NADH_) reduced Ferredoxin (Fd_red_; *ν*_lix, Fd_) and ATP (*ν*_lix, ATP_) molecules needed for the synthesis of the molecule, along with a model of the mechanism used for electron delivery to the microbe [Salimijazi2020b].

For systems that rely upon H_2_-oxidation for electron delivery like the Bionic Leaf [Torella2015a, Liu2016a],

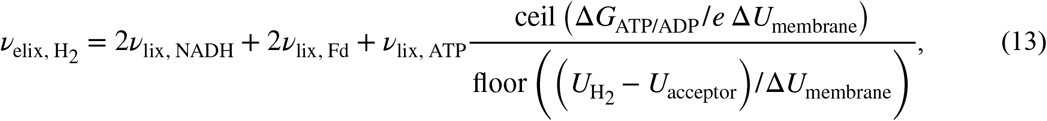

where Δ*G*_ATP/ADP_ is the free energy required for regeneration of ATP, Δ*U*_membrane_ is the potential difference across the cell’s inner membrane due to the proton gradient, *U*_H2_ is the standard potential of proton reduction to H_2_, *U*_acceptor_ is the standard potential of terminal electron acceptor reduction (typically O_2_ + 2*e*^−^ to H_2_O), the ceil function rounds up the nearest integer, and the floor function rounds down to the nearest integer. A full derivation of **Equation 13** can be found in Section 2 (Equations 10 to 20) of the supplement for Salimijazi *et al*. [Salimijazi2020b].

The inner membrane potential difference, Δ*U*_membrane_, is the largest source of uncertainty in this calculation. Therefore, we present a range of efficiency estimates in **Figure 3** and throughout the text for Δ*U*_membrane_ = 80 mV (BioNumber ID (BNID) 10408284 [Milo2010a]) to 270 mV (BNID 107135), with a central value of 140 mV (BNIDs 109774, 103386, and 109775).

For systems that rely upon EEU for electron delivery like *Shewanella oneidensis* [Rowe2021a, Salimijazi2020b],

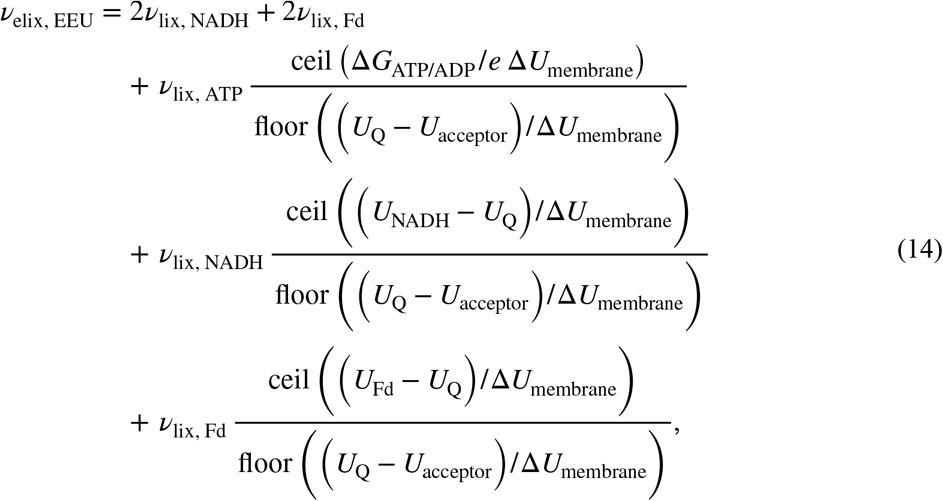

where *U*_Q_ is the redox potential of the inner membrane electron carrier, thought to be ubiquinone [Rowe2018a], *U*_NADH_ is the standard potential of NAD(P)H, and *U*_Fd_ is the standard potential of Ferredoxin. A full derivation of **Equation 14** can be found in Section 7 (Equations 77 to 91) of the supplement for Salimijazi *et al*. [Salimijazi2020b].

The NAD(P)H, ATP and Fd_red_ requirements for lixiviant synthesis were calculated by balancing networks of reactions for the autotrophic synthesis of the molecule from CO_2_ or formate (COOH^−^). We enumerated all reaction steps for the production of 4 environmentally benign lixiviant molecules (acetic, citric, 2,5- diketo-gluconic, and gluconic acid) from acetyl-CoA and using data from the KEGG database in **Table S2** [Kanehisa2000a, Kanehisa2019a, Kanehisa2021a].

Lixiviant synthesis reactions were complemented with reactions for CO_2_-fixation and C1-assimilation. For this article we considered 6 scenarios in which CO_2_ was fixed by the well-known Calvin cycle [Berg2002a], the Reductive Tricarboxylic Acid cycle [Alissandratos2015a, Claassens2016a], Wood-Ljungdahl (WL) Pathway [Berg2002a]; the 3-hydroxypropionate/4-hydroxybutyrate (3HP-4HB) Pathway [BergI2007a, Claassens2016a]; 3-hydroxypropionate (3HP) Cycle [Zarzycki2009a]; and the Dicarboxylate/4-hydroxybutyrate (4HB) Cycle [Huber2008a]. In addition, we also considered the artificial Formolase formate assimilation pathway [Siegel2015a]. These reactions can be found in **Table S3**.

The CO_2_-fixation and C1-assimilation and lixiviants were combined by hand into a set of stoichiometric matrices), **S**_**lix**_, for each reaction network. Stoichiometric matrices are included in **Dataset S1**. Stoichiometric matrices were balanced with a custom flux balance program [Barstow2021b] to find the overall stoichiometry for synthesis of each lixiviant using each CO_2_-fixation or C1-assimilation pathway. The balanced overall stoichiometry for synthesis of each lixiviant by each CO_2_ fixation or C1 assimilation pathway can be found in **Table S4**.

## Results and Discussion

### Mass of Lixiviants Needed for Global Scale CO_2_ Sequestration Can Outstrip Global Supply When De-mineralization Efficiencies Are Low

We plot the mass of lixiviant required for the sequestration of 20 GtCO_2_ per year (the amount of CO_2_ that will need to be sequestered per year in the late 21^st^ century [ipcc2018a]) as a function of the product of the inverse CO_2_ mineralization performance, *ζ*, in **Figure 2**.

**Figure 2.**
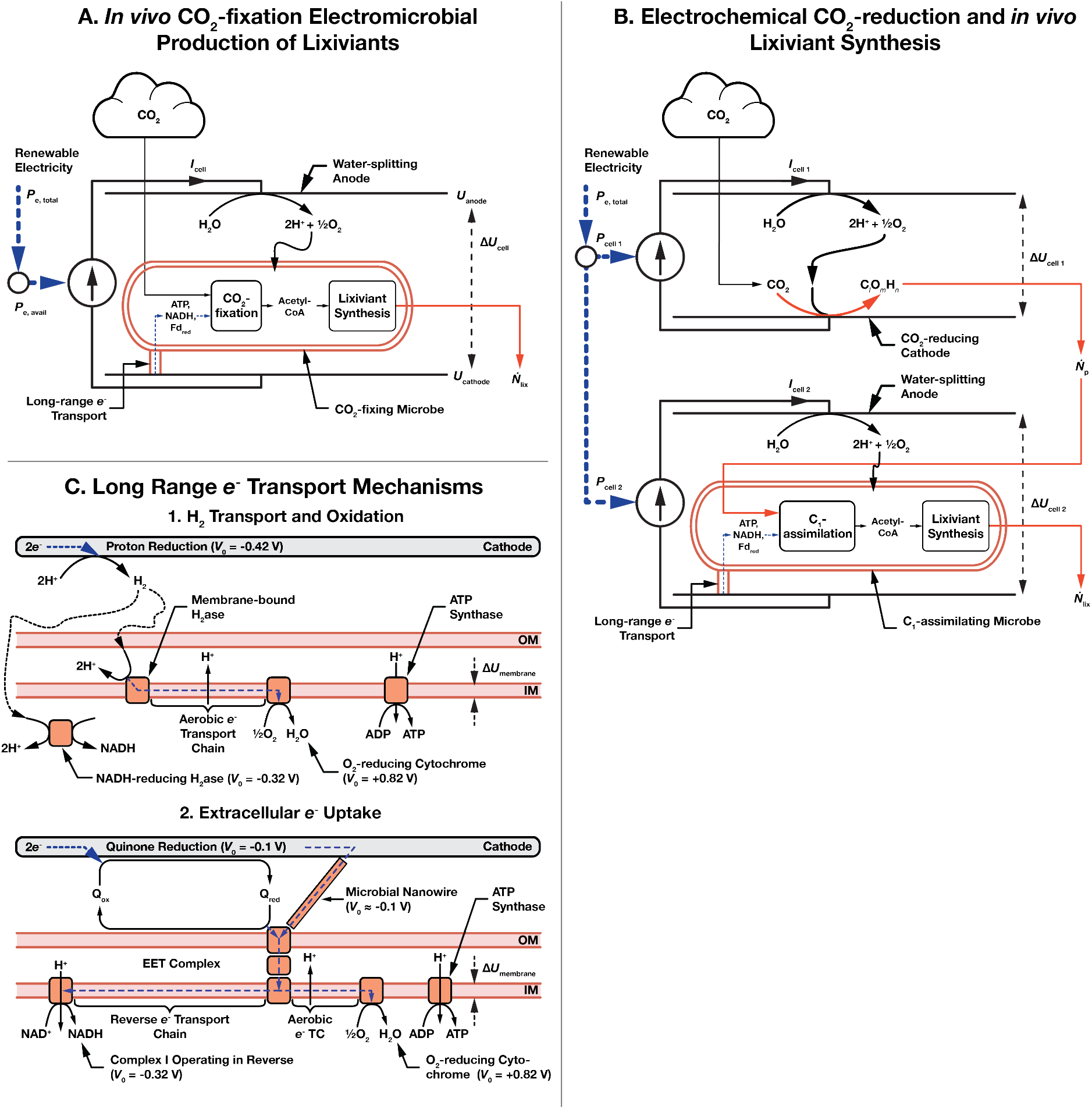
Schematic of electromicrobial production of lixiviants for CO_2_ mineralization. (**A**) Single bio- electrochemical cell system where electricity is used to power *in vivo* CO_2_^−^ and subsequent lixiviant synthesis. (**B**) Dual electrochemical cell system where CO_2_ is reduced in the first cell, and then assimilated in the second cell to produce lixiviant molecules. (**C**) Long range *e*^−^ transfer mechanisms considered in this article. In the first, H_2_ is electrochemically reduced on a cathode, transferred to the microbe by diffusion or stirring, and is enzymatically oxidized. In the second mechanism, extracellular electron uptake (EEU), *e*^−^ are transferred along a microbial nanowire (part of a conductive biofilm), or by a reduced medium potential redox shuttle like a quinone or flavin, and are then oxidized at the cell surface by the extracellular electron transfer (EET) complex. From the thermodynamic perspective considered in this article, these mechanisms are equivalent. Electrons are then transported to the inner membrane where reverse electron transport is used to regenerate NAD(P)H, reduced Ferredoxin (not shown), and ATP [Rowe2018a, Rowe2021a]. Parameters for these systems are shown in **Table 2**.

**Figure 3.**
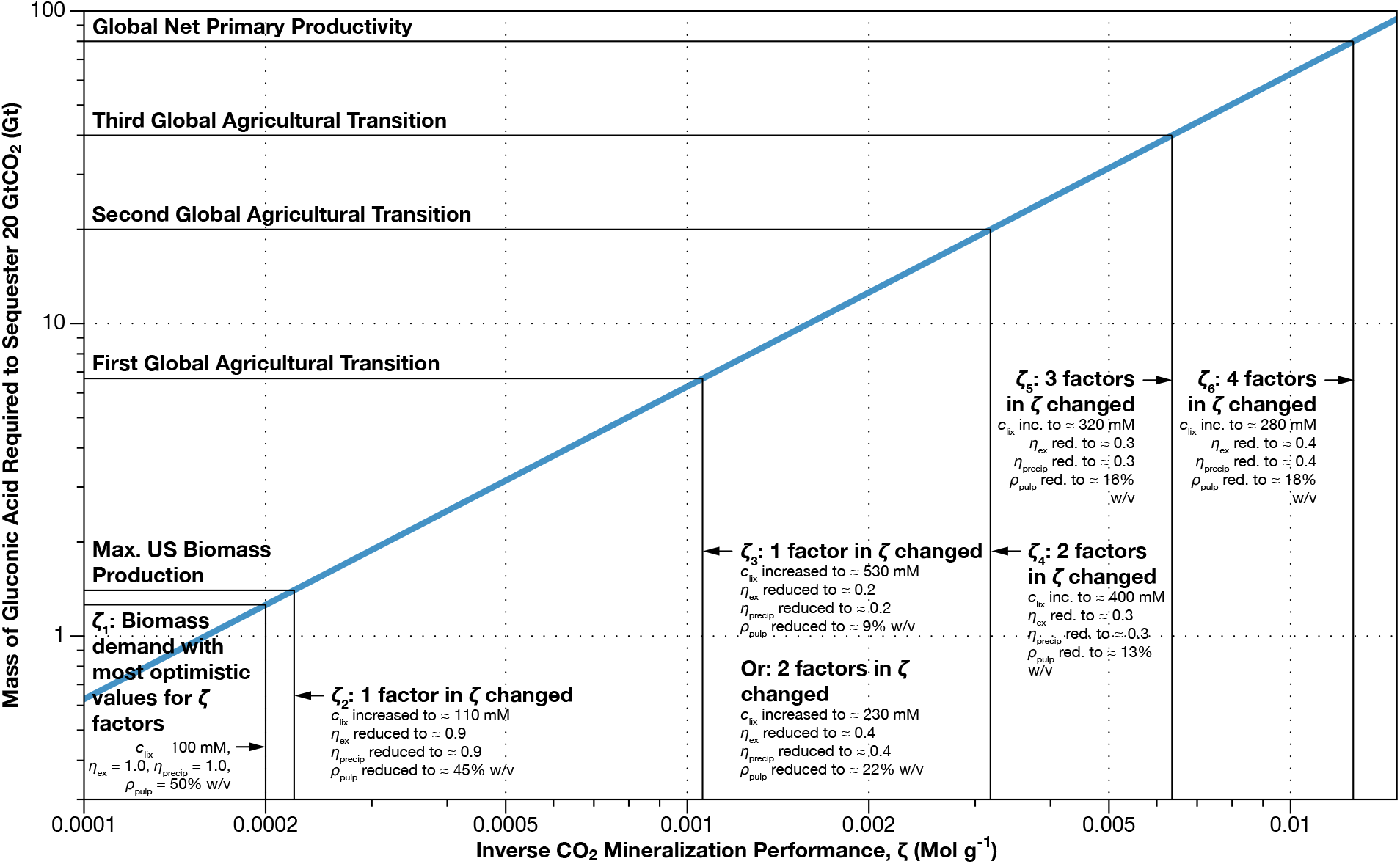
Accelerated mineralization could require hundreds of millions to tens of billions of tonnes of lixiviants per year. If these lixiviants were produced from cellulosic biomass, this could put a significant strain on the world agricultural system. We calculated the mass of lixiviant (*M*_lix_) needed to accelerate the dissolution step of mineralization of 20 GtCO_2_ per year using **Equation 10** as a function of the inverse CO_2_ mineralization performance, *ζ*, the combination of the most uncertain parameters in our estimate of lixiviant mass. We chose to display results for gluconic acid as it has the highest molecular weight and provides an upper bound on the lixiviant mass requirement. Our most optimistic estimate for *ζ* (*ζ*1) is shown as the left most vertical line on the plot. The second marked value of *ζ* (*ζ*_2_) corresponds to a mass of lixiviant equal to all of the cellulosic biomass produced in the United States in a year. The third, fourth and fifth lines (*ζ*_3_ to *ζ*_5_) correspond to increasing biomass withdrawals from the biosphere that come with increasingly severe consequences for agriculture and human society including adoption of vegetation diets, population control and widespread managed agriculture and forestry [Slade2014a]. The sixth (*ζ*_6_) and final line corresponds to the biomass production of the entire world in a year (net primary productivity).This plot can be reproduced with the nlixiviant.py code in the ElectroCO2 repository [Barstow2021b].

What range of values could we expect for the CO_2_ mineralization efficiency? To estimate *ζ* we have made educated guesses for each of the values from the scientific literature. At the optimistic end of the spectrum, we assume that the concentration of lixiviant is 100 mM (corresponding to ≈ pH 2.1 for citric acid, pH 2.4 for gluconic acid, and pH 2.9 for acetic acid; **Note S1**), the extraction and the precipitation efficiency are both 100%, and the pulp density is 50% w/v (500,000 g m^−3^) [MacDonald2007a],

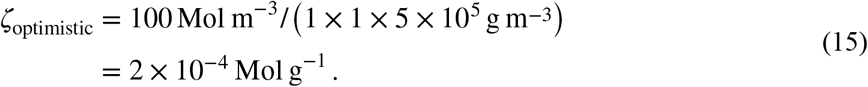

The optimistic value of *ζ* is marked as the furthest left vertical line in **Figure 2**, and corresponds to a consumption of 1.26 Gt of dry lixiviant per year. Even this optimistic scenario corresponds to a significant amount of biomass, accounting for 90% of US biomass production [Perlack2011a] even if cellulosic biomass could be converted to lixiviant with 100% mass conversion efficiency.

**Table 2.**
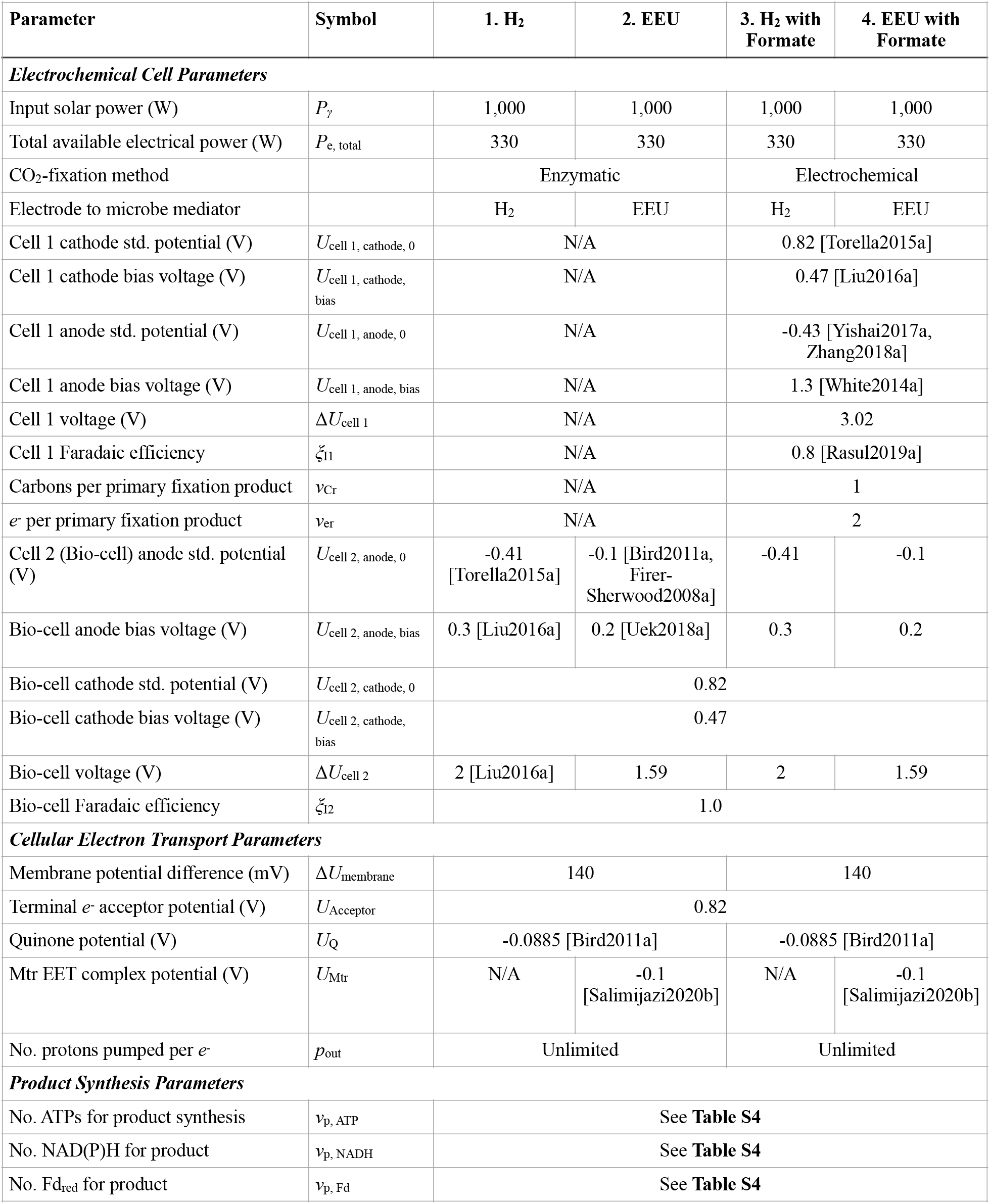
Electromicrobial lixiviant production model parameters. Model parameters used in this article are based upon model parameters used in a previous analysis of the electromicrobial production of the biofuel butanol [Salimijazi2020b]. A sensitivity analysis was performed for all key parameters in this work [Salimijazi2020b].

What are the consequences for lixiviant demand if some of the factors included in *ζ* are slightly less than the optimistic estimates? If just the lixiviant concentration, *c*_lix_, increases by only 10%, or any one of the denominator factors in *ζ* (*η*_ex_, *η*_precip_, *ρ*_pulp_) decreases by 10%, the minimum mass of lixiviant required to sequester 20 GtCO_2_ will rise to 1.4 Gt, equal to the entire biomass production of the United States [Perlack2011a] (**Figure 2**, second vertical line from the left). The same increase in *ζ* can be achieved by a simultaneous 3% increase in *c*_lix_, and 3% reduction in *η*_ex_, *η*_precip_, and *ρ*_pulp_. We have calculated possible combinations of values of *c*_lix_, *η*_ex_, *η*_precip_, and *ρ*_pulp_ that produce each of the values of *ζ* highlighted in **Figure 2** in **Table S5**.

What are the consequences for lixiviant demand if one or more of the factors in *ζ* are significantly less than the optimistic estimates? Slade *et al*. [Slade2014a] calculated the effects of withdrawing increasing quantities of bio-energy from the biosphere. We can make an approximate conversion from bio-energy to dry weight of biomass by dividing by the energy density of dry cellulosic material,

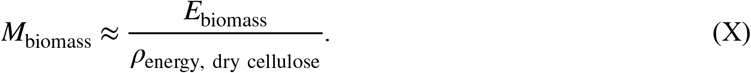

Slade *et al*. [Slade2014a] identified 3 transition points with increasingly restrictive consequences for global civilization (including a combination of crop yield increases, and population, diet and forestry control) that come with increasing biomass use. We have marked these transition points as the third, fourth and fifth horizontal lines from the bottom of **Figure 2**. We have marked values of *ζ* that correspond to these transition points as the third, fourth and fifth vertical lines from the left in **Figure 2**.

A significant change in one of the factors of *ζ* or two smaller simultaneous changes is required for lixiviant demand to pass the first consequential transition identified by Slade *et al*. [Slade2014a]. The first transition occurs when the withdrawal of bio-energy from the biosphere exceeds 100 exajoules per year (EJ) (corresponding to ≈ 7 Gt of dry biomass). Exceeding this withdrawal rate will require that crop yields keep pace with demand; and either adoption of vegetarian diets, or a low global population (< 9 billion), or limited deforestation. Increasing the lixiviant demand rate to ≈ 7 Gt per year occurs when ζ rises to 1 × 10^−6^ Mol g^−1^. This increase in *ζ* will happen if *c*_lix_ rises by a factor of ≈ 5 to 530 mM, or a reduction in any one of the of the denominator factors (*η*_ex_, *η*_precip_, and *ρ*_pulp_) to ≈ 1/5^th^ of its optimistic value (**Figure 2, Table S5**). *ζ* can also rise to 10^−6^ Mol g^−1^ if *c*_lix_ rises by a factor of ≈ 2, and one of the denominator factors falls to ≈ 1/2 of its optimistic value, or two of the the denominator factors fall to ≈ 1/2 of their optimistic value. Alternatively, the same increase in *ζ* can also happen if *c*lix increases by ≈ 50% (3/2), and the denominator factors all decrease to about 2/3^rds^ of their optimistic values (**Table S5**).

Significant changes in two factors contributing to *ζ* are required for lixiviant demand to pass the second consequential transition identified by Slade *et al*. [Slade2014a]. This second transition occurs when the withdrawal of bio-energy from the biosphere exceeds 300 EJ per year (≈ 20 Gt of dry biomass per year). Exceeding this withdrawal rate will require that increases in crop yields outpace demand; and either adoption of vegetarian diets, a low population or limited deforestation. Increasing the lixiviant demand rate to 20 Gt occurs if there are simultaneous reductions in two of the three denominator factors of *ζ* to ≈ 1/4^th^ of their optimistic value, or an increase in *c*_lix_ to ≈ 400 mM (a factor of 4) (**Figure 2** and **Table S5**). Alternatively, a doubling of *c*_lix_ to ≈ 200 mM, and a reduction in all the denominator factors to 1/2 their optimistic value will also raise lixiviant demand to 20 Gt (**Table S5**).

Significant changes in three factors contributing to *ζ* are required for lixiviant demand to pass the third consequential transition identified by Slade *et al*. [Slade2014a]. The third transition point occurs when bio-energy withdrawal exceeds 600 EJ yr^−1^ (≈ 40 Gt of dry biomass per year). Exceeding this withdrawal rate requires high input farming, high increases in crop yields, limiting global population to < 9 billion, and adoption of either vegetarian diets or managed forestry [Slade2014a]. Increasing the lixiviant demand rate to 40 Gt can occur if *c*lix triples to 300 mM, and 2 of the denominator factors are reduced to ≈ 1/3^rd^ of their optimistic values (**Figure 2** and **Table S5**).

Finally, the lixiviant demand rate can thoroughly bust the Earth’s biomass budget, exceeding net primary productivity (NPP) of 120 EJ yr^−1^ (80 Gt dry biomass) if *c*lix increases to 280 mM, and all 3 denominator factors are reduced to ≈ 1/3^rd^ of their optimistic values (**Figure 2** and **Table S5**).

Taken together, the results presented here suggest that CO_2_ mineralization accelerated with biologically produced lixiviants could (although this is definitely not guaranteed) place an undesirable burden on the Earth’s biosphere.

### Electromicrobial Production Could Produce Lixiviants at a Cost of a Few Hundred Dollars per Tonne

Electromicrobial production technologies already have lab scale efficiencies that can exceed the theoretical upper limit efficiencies of most forms of photosynthesis [Torella2015a, Liu2016a, Haas2018a], and have even further room to improve [Salimijazi2020b, Wise2021a]. This means that electromicrobial production might be able to produce lixiviants for CO_2_ mineralization from electricity and CO_2_ without needing to compete for land with agriculture and wilderness.

We used our theory of electromicrobial production (**Theory** and refs [Salimijazi2020b, Wise2021a]) to calculate the minimum electricity needs, and hence minimum solar electricity costs needed to produce a tonne of 4 different lixiviant compounds: acetic acid, citric acid, 2,5-diketogluconic acid, and gluconic acid (**Figure 3)**.

The most expensive lixiviant to synthesize is acetic acid produced with the 4HB CO_2_-fixation pathway and with electrons supplied with extracellular electron uptake (EEU) at a cost of 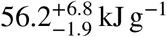. Assuming that the US Department of Energy’s solar PV electricity price projection for 2030 of 3 ¢ per kilowatt-hour can be achieved, this translates to a cost of $468 per tonne of acetic acid (right hand side axes in **Figure 3**).

As in our earlier analyses [Salimijazi2020b, Wise2021a] modifying the CO_2_ fixation method from the least efficient (the 4HB pathway) to the most efficient (the Wood-Ljungdahl pathway) can reduce energy costs of electromicrobial production by almost a factor of 2 [Salimijazi2020b, Wise2021a]. Likewise, switching the electron delivery mechanism to H_2_-oxidation further reduces energy costs of production. The lowest cost method for producing acetic acid is with the Wood-Ljungdahl CO_2_-fixation pathway and with electrons supplied by H_2_-oxidation, which results in a cost of 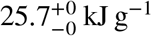, or $214 per tonne. The lowest cost lixiviant is citric acid, with a minimum cost of 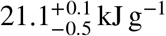 ($175 per tonne) when produced with the Wood-Ljungdahl pathway and with electron delivery by H_2_-oxidation.

Electromicrobial lixiviant production is more expensive than biomass production, even with projected 2030 solar PV prices, but might still achieve cost parity. The farm gate cost of cellulosic biomass ranges from $39.7/dry tonne for loblolly pine wood chip to $72.3/dry tonne for switchgrass [Liu2015a], between 3 and 10 times cheaper than electromicrobially produced lixiviants. However, these costs do not include the cost of conversion of cellulosic biomass to a lixiviant. It is estimated that the production cost of cellulosic ethanol is $2.65 per US gallon ($890 per tonne), and it is reasonable to assume that lixiviant production would incur similar costs. Electromicrobial production of lixiviants could still achieve cost parity with biomass-derived lixiviants by directly producing the lixiviant and avoiding conversion costs.

### Electromicrobially Produced Lixiviants Might Enable Cost-competitive CO_2_ Mineralization

The costs of CO_2_ mineralization with electromicrobially produced lixiviants are high, but could still enable cost effective CO_2_ mineralization. We have plotted the amount of energy needed to synthesize enough acetic, gluconic, citric and 2,5-diketo-gluconic acid to sequester 1 tonne of CO_2_ as a function of the inverse CO_2_ mineralization economy, *ζ*, in **Figure 4**. While acetic acid is the most expensive lixiviant to produce on a per tonne basis, for a given value of *ζ*, it produces the lowest cost CO_2_ mineralization.

**Figure 4.**
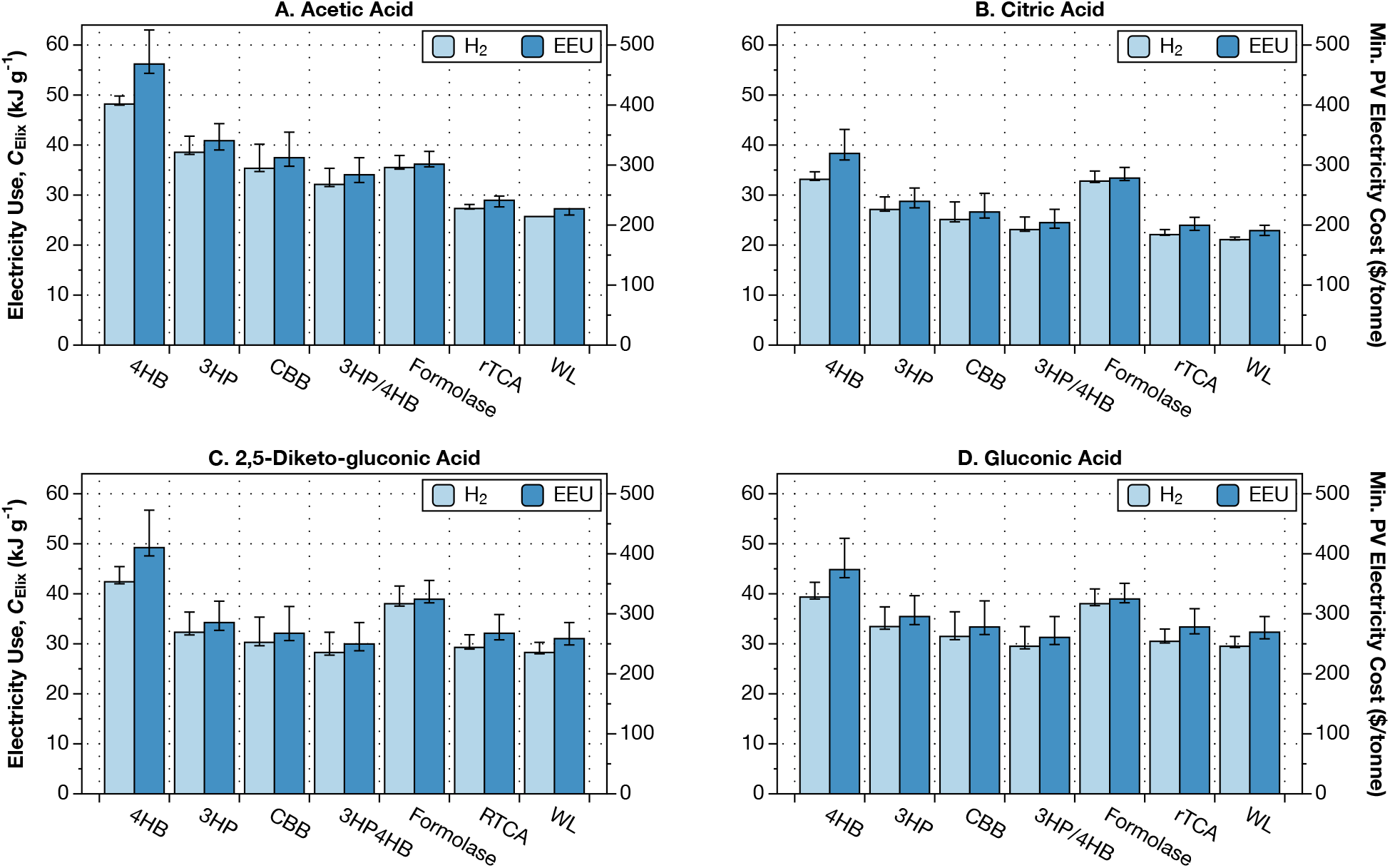
Electromicrobial production technology could reduce the electrical energy costs of lixiviant production to a few tens of kilojoules per gram. Energy and financial costs for producing 4 lixiviant molecules are shown in each panel: (**A**) acetic acid, (**B**) citric acid, (**C**) 2,5-diketo-gluconic acid (DKG), and (**D**) gluconic acid. The electrical energy cost of producing a gram of each lixiviant is shown the left hand side *y*-axis for each sub-plot. The dollar cost of producing a tonne of the lixiviant using electricity supplied by solar photovoltaics at a cost of 3¢ per kWh (the US Department of Energy’s cost target for solar electricity for 2030 [SunShot]). This plot can be reproduced using the efficiency.py code in the ElectroCO2 repository [Barstow2021b].

**Figure 5.**
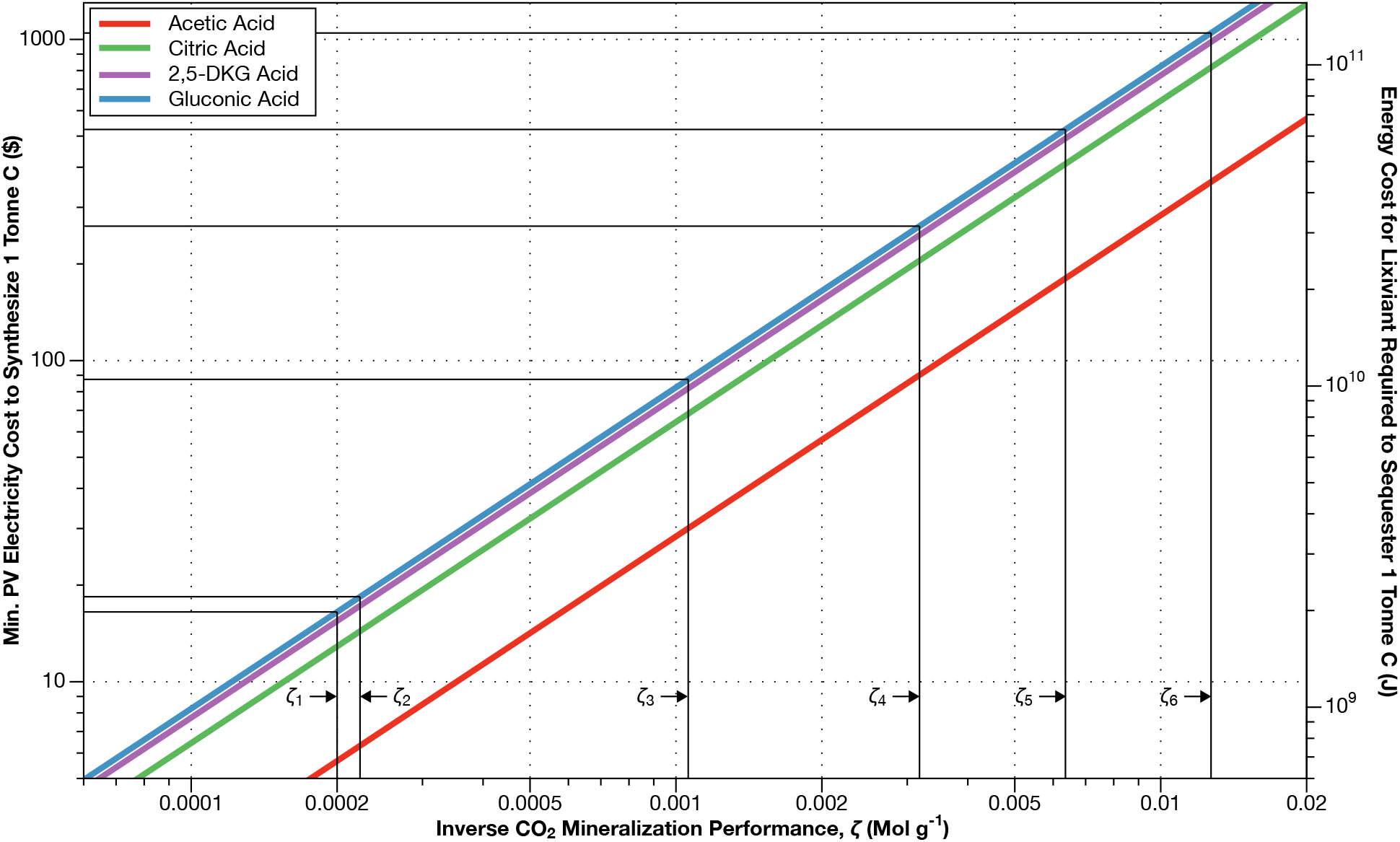
Electromicrobial production technology could enable production of enough lixiviants to sequester 1 tonne of CO_2_ for less than $100. We combined our lixiviant mass requirements from **Figure 3**, with our estimates for the energy and financial cost of producing a tonne of each lixiviant compound with H_2_-mediated EMP using CO_2_-fixation with the Calvin cycle (basically the Bionic Leaf configuration [Torella2015a, Liu2016a]) from **Figure 4**. For illustrative purposes we have marked the values of the inverse CO_2_ mineralization performance (*ζ*_1_ to *ζ*_6_) highlighted in **Figure 3**, and the corresponding cost to sequester a tonne of CO_2_ as an intersecting horizontal line. However, it is important to note that in this case, no cellulosic biomass is produced. This plot can be reproduced using the Clixiviant.py code in the ElectroCO2 repository [Barstow2021b].

For the most optimistic value of *ζ* (2 × 10^−4^ Mol g^−1^), the cost of electricity (at projected 2030 PV prices) needed to make enough gluconic acid to sequester 1 tonne of CO_2_ is $17 (and only $6 for acetic acid) (**Figure 4**). Even when *ζ* rises to 1 × 10-3 Mol g^−1^ (corresponding to a biomass drain from the biosphere that would prompt significant changes to global agriculture) the cost of sequestering a tonne of CO_2_ only rises to $87 when using gluconic acid, and $30 when using acetic acid (**Figure 4**).

## Conclusions

CO_2_ sequestration at the scale discussed in this article (20 GtCO2 yr^−1^) is not likely to be needed for approximately 50 years from the time of writing (around 2070). This means that there is time to identify technologies that could meet this need and refine them to do it. Weathering of ultramafic rocks and subsequent mineralization of CO_2_ almost certainly has the capacity to deal with the excess CO_2_ in the atmosphere, but accelerating this process remains a challenge.

Accelerating the weathering of ultramafic materials to the rate necessary to keep climate change within acceptable limits with organic lixiviants made from cellulosic biomass has the potential to monopolize the world’s biomass supply. Even under the most optimistic estimate of CO_2_ mineralization performance, sequestration of 20 GtCO_2_ per year could use 90% of the biomass production of the entire United States (**Figure 2**). If the CO_2_ mineralization performance were to slip slightly, accelerated CO_2_ mineralization could force undesirable changes to the world agricultural system and society (**Figure 2**).

Electromicrobial production of organic lixiviants could enable accelerated CO_2_ mineralization without competing for agricultural land. While EMP technologies only exist in the lab today, their high lab-scale and even higher predicted solar to product conversion efficiencies mean that they could be an effective tool in CO2 management. In this article, we demonstrate that organic lixiviants can be produced by EMP at the cost of ≈ $200 to $400 per tonne assuming solar electricity is supplied at a cost of 3¢ per kWh (a target for 2030 solar electricity costs set by the US Department of Energy [SunShot]) (**Figure 3**).

Electromicrobially produced lixiviants could enable large scale CO_2_ mineralization at low costs. We show that even with modest CO_2_ mineralization performance, the cost of making the lixiviants needed to sequester a tonne of CO_2_ could be kept below $100 per tonne, even with 2030 solar electricity costs (**Figure 4**). It is highly likely that many more halvings of solar electricity cost will occur between 2030 and 2070, further reducing the cost of CO_2_ mineralization.

What’s the best way to achieve the potential of EMP for CO_2_ mineralization? Until recently, the difficulty of adding CO_2_ fixation to a non-CO_2_-fixing organism; uncertainty about the efficiency of electron uptake by EET and even if it can reduce the NADH needed for CO_2_ fixation; and the difficulty of engineering non-model organisms like *G. oxydans* has made a project like this look unfeasible. However, recently Rowe *et al*. [Rowe2018a] discovered that *S. oneidensis* can use imported electrons to reduce NADH Meanwhile, Gleizer *et al*. [Gleizer2019a] transformed the lab workhouse microbe *E. coli* to fix CO_2_.

Furthermore, we have recently discovered the genes that code for this electron uptake pathway in the electroactive microbe in *S. oneidensis*; and have used the Knockout Sudoku technology [Baym2016a, Anzai2017a] to build a whole genome knockout collection of the mineral-dissolving microbe *G. oxydans*, the first step in whole genome engineering. Added together these breakthroughs make something that appeared almost impossible a year ago look tantalizingly close.

## End Notes

### Code Availability

All code used in calculations in this article is available at https://github.com/barstowlab/electroCO2 and is archived on Zenodo [Barstow2021b].

### Materials & Correspondence

Correspondence and material requests should be addressed to B.B.

### Author Contributions

Conceptualization, B.B.; Methodology, B.B. and S.M.; Investigation, S.M., A.D., L.G., L.L., K.S., I.T., J.Z., and B.B; Writing - Original Draft, A.D., L.G., L.L., K.S., I.T., J.Z., and B.B.; Writing - Review and Editing, B.B.; Resources, B.B.; Supervision, B.B..

## Acknowledgements

This work was supported by Cornell University startup funds, a Career Award at the Scientific Interface from the Burroughs Welcome Fund, ARPA-E award DE-AR0001341, and by U.S. Department of Energy Biological and Environmental Research grant DE-SC0020179 to B.B..

