## Supplementary material for "Practical and Thermodynamic Constraints on Electromicrobially-Accelerated CO_2_ Mineralization": SI Text

### Supplementary Information Tables

**Table S1.** Molecular weights and energy densities for lixiviant molecules considered in this article.

**Table S2.** Reactions for synthesis of lixiviant molecules considered in this article.

**Table S3.** Carbon-fixation and -assimilation reactions considered in this article.

**Table S4.** Net molecular input requirements for lixiviant synthesis by 6 naturally-occurring $CO_2$-fixation cycles and the synthetic Formolase formate assimilation pathway.

### Supplementary Notes

**Note S1**. Calculation of biolixiviant pH.

### Supplementary Information Datasets

**Dataset S1.** Enzymatic reactions for synthesis of lixiviant compounds from $CO_2$ or formic acid.

| Lixiviant Compound | Molecular Weight (Da) | Molecular Formula | Carbons per Molecule |
|---|---|---|---|
| Acetic Acid | 60.052 | $CH_3COOH$ | 2 |
| Citric Acid | 192.124 | $C_6H_8O_7$ | 6 |
| 2,5-Diketo-Gluconic Acid | 191.12 | $C_6H_8O_7$ | 6 |
| Gluconic Acid | 196.16 | $C_6H_{12}O_7$ | 6 |
| Glucose | 180.16 | $C_6H_{12}O_6$ | 6 |

**Table S1.** Molecular weights and energy densities for lixiviant molecules considered in this article.

| Reaction | Reference |
|---|---|
| ***1. Acetic Acid*** | |
| Acetyl-CoA + ADP + phosphate → Acetate + ATP + CoA | KEGG R0229 |
| ***2. Citric Acid*** | |
| Pyruvate + $CO_2$ → Oxaloacetate | KEGG RC00040 |
| Acetyl-CoA + $H_2O$ + Oxaloacetate → Citrate + CoA | KEGG RC00351 |
| ***3. 2,5-Diketo-Gluconic Acid*** | |
| ATP + Pyruvate + $HCO_3^-$ → Orthophosphate + Oxaloacetate | KEGG R00344 |
| ATP + Oxaloacetate → Phosphoenolpyruvate + $CO_2$ | KEGG R00341 |
| Phosphoenolpyruvate + $H_2O$ → 2-Phospho-D-glycerate | KEGG R00658 |
| 2-Phospho-D-glycerate → 3-Phospho-D-glycerate | KEGG R01518 |
| ATP + 3-Phospho-D-glycerate → 3-Phospho-D-glyceroyl phosphate | KEGG R01512 |
| 3-Phospho-D-glyceroyl phosphate + NADH + $H^+$ → D-Glyceraldehyde 3-phosphate + Orthophosphate | KEGG R01061 |
| D-Glyceraldehyde 3-phosphate → Glycerone phosphate | KEGG R01015 |
| Glycerone phosphate + D-Glyceraldehyde 3-phosphate → D-Fructose 1,6-bisphosphate | KEGG R01068 |
| D-Fructose 1,6-bisphosphate + $H_2O$ → D-Fructose 6-phosphate + Orthophosphate | KEGG R00762 |
| D-Fructose 6-phosphate → D-Glucose 6-phosphate | KEGG R00771 |
| D-Glucose 6-phosphate + $H_2O$ → D-Glucose + Orthophosphate | KEGG R00303 |
| beta-D-Glucose → D-Glucono-1,5-lactone + NADH+ $H^+$ | KEGG R01521 |
| D-Glucono-1,5-lactone + $H_2O$ → D-Gluconate | KEGG R01519 |
| D-Gluconate → 2-Keto-D-gluconic acid + $H^+$ | KEGG R01739 |
| 2-Keto-D-gluconic acid → 2,5-Diketo-D-gluconic acid+ $H^+$ + NADPH | KEGG R05823 |
| ***4. Gluconic Acid*** | |
| ATP + Pyruvate + $HCO_3^-$ → ADP + Phosphate + Oxaloacetate | KEGG R00344 |
| ATP + Oxaloacetate → ADP + Phosphoenolpyruvate + $CO_2$ | KEGG R00341 |
| Phosphoenolpyruvate + $H_2O$ → 2-phospho-D-glycerate | KEGG R00658 |
| 2-phospho-D-glycerate → 3-phospho-D-glycerate | KEGG R01518 |
| ATP + 3-phospho-D-glycerate → ADP + 3-phospho-D-glyceroyl phosphate | KEGG R01512 |
| 3-phospho-D-glyceroyl phosphate + NADH + $H^+$ → D-glyceraldehyde 3-phosphate + Phosphate + $NAD^+$ | KEGG R01061 |
| D-glyceraldehyde 3-phosphate → Glycerone phosphate | KEGG R01015 |
| Glycerone phosphate + D-glyceraldehyde 3-phosphate → D-fructose 1,6-bisphosphate | KEGG R01068 |
| D-fructose 1,6-bisphosphate + $H_2O$ → D-fructose 6-phosphate + Phosphate | KEGG R00762 |
| D-fructose 6-phosphate → D-glucose 6-phosphate | KEGG R00771 |
| D-glucose 6-phosphate + $H_2O$ → D-glucose + Phosphate | KEGG R00303 |
| D-glucose + $NAD(P)^+$ → D-glucono-1,5-lactone + NAD(P)H + $H^+$ | KEGG R01520 |
| D-glucono-1,5-lactone + $H_2O$ → D-gluconate | KEGG R01519 |

**Table S2.** Reactions for synthesis of lixiviant molecules considered in this article. Reactions for lixiviant synthesis from acetyl-CoA, NAD(P)H, Ferredoxin and ATP were assembled from data from the KEGG database [Kanehisa2000a, Kanehisa2019a, Kanehisa2021a]

| **Reaction** | **Reference** |
|---|---|
| ***1. Calvin Cycle (CBB)*** | |
| 2 $CO_2$ + 7 ATP + 4 NADH → 1 Acetyl-CoA | Salimijazi *et al*. [Salimijazi2020b]. |
| 3 $CO_2$ + 7 ATP + 5 NADH → 1 Pyruvate | Salimijazi *et al* [Salimijazi2020b]. |
| ***2. Wood-Ljungdahl Pathway (WL)*** | |
| 4 $CO_2$ + 2 ATP + 8 NADH → 2 Acetyl-CoA | Berg [Berg2011a]. |
| 2 $Fd_{red}$ + Acetyl-CoA + $CO_2$ → Pyruvate | KEGG R01196. |
| ***3. Reductive TCA Cycle (RTCA)*** | |
| 4 CO2 + 4 ATP + 8 NADH → 2 Acetyl-CoA | Alissandratos *et al.* [Alissandratos2015a], Claassens *et al*. [Claassens2016a]. |
| 2 $Fd_{red}$ + Acetyl-CoA + $CO_2$ → Pyruvate | KEGG R01196. |
| ***4. 3-hydroxypropionate/4-hydroxybutyrate Cycle (3HP4HB)*** | |
| 6 $HCO_3^-$ + 10 ATP + 10 NADH → 2 pyruvate | Berg *et al*. [BergI2007a], Claassens *et al*. [Claassens2016a]. |
| 2 Pyruvate → 2 Acetyl-CoA + 2 NADH + 2 $CO_2$ | Berg [Berg2002a], Schomburg *et al.* [Schomburg2017a]. |
| ***5. 3-hydroxypropionate Cycle (3HP)*** | |
| 6 $HCO_3^-$ + 10 ATP + 12 NADH → 2 Pyruvate | Zarzycki *et al*. [Zarzycki2009a], Herter *et al*. [Herter2002a], Berg [Berg2002a]. |
| 2 Pyruvate → 2 Acetyl-CoA + 2 NADH + 2 $CO_2$ | Zarzycki *et al*. [Zarzycki2009a], Herter *et al*. [Herter2002a], Berg [Berg2002a]. |
| ***6. 4-hydroxybutyrate Cycle (4HB)*** | |
| 1 $CO_2$ + 1 $HCO_3^-$ + 3 ATP + 1 NADH + 6 $Fd_{red}$ → 1 Acetyl-CoA | Huber *et al*. [Huber2008a]. |
| 2 Pyruvate → 2 Acetyl-CoA + 2 NADH + 2 $CO_2$ | Berg [Berg2002a], Schomburg *et al.* [Schomburg2017a]. |
| ***7. Formolase Pathway (FORM)*** | |
| 6 $HCO_2^-$ + 10 ATP + 4 NADH → 2 3-PG | Siegel *et al*. [Siegel2015a], Bar-Even *et al*. [Bar-Even2016a]. |
| 2 3-PG → 2 Pyruvate + 2 ATP | Berg [Berg2002a]. |
| 2 Pyruvate → 2 Acetyl-CoA + 2 NADH + 2 $CO_2$ | Berg [Berg2002a], Schomburg *et al.* [Schomburg2017a]. |

**Table S3.** $CO_2$-fixation and $C_1$-assimilation reactions. $CO_2$-fixation and $C_1$-assimilation reactions considered in this article were first assembled in Salimijazi *et al*. [Salimijazi2020b] and are restated here for convenience. Overall reactions for production of metabolic intermediates by 6 naturally-occurring $CO_2$-fixation cycles and the synthetic Formolase formate assimilation pathway, and the FeMoCo nitrogenase $N_2$-fixation reaction. Reactions can be referenced KEGG database [Kanehisa2000a, Kanehisa2019a, Kanehisa2021a]. $Fd_{red}$: Reduced Ferredoxin; 3-PG: 3-Phosphoglycerate.

| **Scenario** | **ATP** | **NAD(P)H** | **$Fd_{red}$** | **$CO_2$** | **$HCO_3^-$** | **$HCO_2^-$** | **Total C** | **Target Molecule** | **Target Formula** | **Target** |
|---|---|---|---|---|---|---|---|---|---|---|
| Acetic_3HP | 4 | 5 | 0 | -1 | 3 | 0 | 2 | Acetate | $CH_3COOH$ | 1.0 |
| Acetic_3HP4HB | 4 | 4 | 0 | -1 | 3 | 0 | 2 | Acetate | $CH_3COOH$ | 1.0 |
| Acetic_4HB | 2 | 1 | 6 | 1 | 1 | 0 | 2 | Acetate | $CH_3COOH$ | 1.0 |
| Acetic_CBB | 6 | 4 | 0 | 2 | 0 | 0 | 2 | Acetate | $CH_3COOH$ | 1.0 |
| Acetic_FORM | 3 | 1 | 0 | -1 | 0 | 3 | 2 | Acetate | $CH_3COOH$ | 1.0 |
| Acetic_RTCA | 1 | 4 | -0 | 2 | 0 | 0 | 2 | Acetate | $CH_3COOH$ | 1.0 |
| Acetic_WL | -0 | 4 | -0 | 2 | 0 | 0 | 2 | Acetate | $CH_3COOH$ | 1.0 |
| Citric_3HP | 10 | 11 | 0 | 0 | 6 | 0 | 6 | Citrate | $C_6H_8O_7$ | 1.0 |
| Citric_3HP4HB | 10 | 9 | 0 | 0 | 6 | 0 | 6 | Citrate | $C_6H_8O_7$ | 1.0 |
| Citric_4HB | 6 | 3 | 12 | 4 | 2 | 0 | 6 | Citrate | $C_6H_8O_7$ | 1.0 |
| Citric_CBB | 14 | 9 | 0 | 6 | 0 | 0 | 6 | Citrate | $C_6H_8O_7$ | 1.0 |
| Citric_FORM | 8 | 3 | 0 | 0 | 0 | 6 | 6 | Citrate | $C_6H_8O_7$ | 1.0 |
| Citric_RTCA | 4 | 8 | 2 | 6 | 0 | 0 | 6 | Citrate | $C_6H_8O_7$ | 1.0 |
| Citric_WL | 2 | 8 | 2 | 6 | 0 | 0 | 6 | Citrate | $C_6H_8O_7$ | 1.0 |
| DKG_3HP | 16 | 12 | 0 | -2 | 8 | 0 | 6 | 2 5-DKG | $C_6H_8O_7$ | 1.0 |
| DKG_3HP4HB | 16 | 10 | 0 | -2 | 8 | 0 | 6 | 2 5-DKG | $C_6H_8O_7$ | 1.0 |
| DKG_4HB | 12 | 2 | 16 | 2 | 4 | 0 | 6 | 2 5-DKG | $C_6H_8O_7$ | 1.0 |
| DKG_CBB | 20 | 10 | 0 | 4 | 2 | 0 | 6 | 2 5-DKG | $C_6H_8O_7$ | 1.0 |
| DKG_FORM | 14 | 4 | 0 | -2 | 2 | 6 | 6 | 2 5-DKG | $C_6H_8O_7$ | 1.0 |
| DKG_RTCA | 10 | 8 | 4 | 4 | 2 | 0 | 6 | 2 5-DKG | $C_6H_8O_7$ | 1.0 |
| DKG_WL | 8 | 8 | 4 | 4 | 2 | 0 | 6 | 2 5-DKG | $C_6H_8O_7$ | 1.0 |
| Gluconate_3HP | 16 | 13 | 0 | -2 | 8 | 0 | 6 | D-Gluconate | $C_6H_{12}O_7$ | 1.0 |
| Gluconate_3HP4HB | 16 | 11 | 0 | -2 | 8 | 0 | 6 | D-Gluconate | $C_6H_{12}O_7$ | 1.0 |
| Gluconate_4HB | 12 | 5 | 12 | 2 | 4 | 0 | 6 | D-Gluconate | $C_6H_{12}O_7$ | 1.0 |
| Gluconate_CBB | 20 | 11 | 0 | 4 | 2 | 0 | 6 | D-Gluconate | $C_6H_{12}O_7$ | 1.0 |
| Gluconate_FORM | 12 | 5 | 0 | 0 | 0 | 6 | 6 | D-Gluconate | $C_6H_{12}O_7$ | 1.0 |
| Gluconate_RTCA | 10 | 9 | 4 | 4 | 2 | 0 | 6 | D-Gluconate | $C_6H_{12}O_7$ | 1.0 |
| Gluconate_WL | 8 | 9 | 4 | 4 | 2 | 0 | 6 | D-Gluconate | $C_6H_{12}O_7$ | 1.0 |

**Table S4.** Net molecular input requirements for lixiviant synthesis by 6 naturally-occurring $CO_2$-fixation cycles and the synthetic Formolase formate assimilation pathway. $Fd_{red}$: Reduced Ferredoxin. 3HP: 3-hydroxypropionate cycle [Zarzycki2009a]; 3HP4HB: 3-hydroxypropionate/4-hydroxybutyrate pathway [BergI2007a, Claassens2016a]; 4HB: Dicarboxylate/4-hydroxybutyrate cycle [Huber2008a]; Calvin-Benson-Bassham cycle [Berg2002a]; FORM: Formolase formate assimilation pathway [Siegel2015a]; RTCA: Reductive Tricarboxylic Acid cycle [Alissandratos2015a, Claassens2016a]; WL: Wood-Ljungdahl (WL) Pathway [Berg2002a]. Results can be reproduced running the BALANCE.PY code in the ELECTROCO2 repository [Barstow2021b].

| # | Inverse $CO_2$ Mineralization Economy, $\zeta$ (Mol $g^{-1}$) | Lixiviant Concentration, $c_{lix}$ (Moles per $m^3$ or mM) | Extraction Efficiency, $\eta_{ex}$ | Precipitation Efficiency, $\eta_{precip}$ | Pulp Density, $\rho_{pulp}$ (grams per $m^3$) | Note |
|---|---|---|---|---|---|---|
| $\zeta_1$ | $2.00 \times 10^{-4}$ | 100.0 | 1.00 | 1.00 | $5.00 \times 10^5$ | Our most optimistic estimate of inverse $CO_2$ mineralization economy, $\zeta$. |
| $\zeta_2$ | $2.23 \times 10^{-4}$ | 100.0 | 1.00 | 0.90 | $5.00 \times 10^5$ | Value of $\zeta$ corresponding to use of entire US biomass production. |
| | | 111.5 | 1.00 | 1.00 | $5.00 \times 10^5$ | |
| | | 100.0 | 0.90 | 1.00 | $5.00 \times 10^5$ | |
| | | 102.8 | 0.97 | 0.97 | $4.87 \times 10^5$ | |
| $\zeta_3$ | $1.06 \times 10^{-3}$ | 100.0 | 1.00 | 0.19 | $5.00 \times 10^5$ | Value of $\zeta$ corresponding to first global agricultural transition identified by Slade *et al.* [Slade2014a]. |
| | | 100.0 | 0.19 | 1.00 | $5.00 \times 10^5$ | |
| | | 530.0 | 1.00 | 1.00 | $5.00 \times 10^5$ | |
| | | 100.0 | 1.00 | 1.00 | $9.40 \times 10^4$ | |
| | | 100.0 | 0.44 | 0.44 | $5.00 \times 10^5$ | |
| | | 229.9 | 1.00 | 1.00 | $2.18\times 10^5$ | |
| | | 151.7 | 0.66 | 0.66 | $3.30\times 10^5$ | |
| $\zeta_4$ | $3.18 \times 10^{-3}$ | 100.0 | 0.25 | 0.25 | $5.00 \times 10^5$ | Value of $\zeta$ corresponding to second global agricultural transition identified by Slade *et al.* [Slade2014a]. |
| | | 398.9 | 1.00 | 1.00 | $1.25 \times 10^5$ | |
| | | 199.8 | 0.50 | 0.50 | $2.50 \times 10^5$ | |
| $\zeta_5$ | $6.36 \times 10^{-3}$ | 237.5 | 0.42 | 0.42 | $2.10 \times 10^5$ | Value of $\zeta$ corresponding to third global agricultural transition identified by Slade *et al.* [Slade2014a]. |
| | | 316.8 | 0.32 | 0.32 | $5.00 \times 10^5$ | |
| | | 316.8 | 1.00 | 0.32 | $1.58 \times 10^5$ | |
| $\zeta_6$ | $1.27 \times 10^{-2}$ | 282.5 | 0.35 | 0.35 | $1.77 \times 10^5$ | Value of $\zeta$ corresponding to use of entire global net primary production [Slade2014a]. |

**Table S5.** Possible combinations factors that produce significant values of inverse $CO_2$ mineralization economy, $\zeta$. We calculated possible combinations of values of $c_{lix}$, $\eta_{ex}$, $\eta_{precip}$, and $\rho_{pulp}$ that produce each of the values of $\zeta$ highlighted in **Figure 2**.

### Supplementary Note 1: Calculation of Lixiviant pH

What is the pH of a weak organic acid, with a given $pK_a$, at a particular analytical concentration? We have adapted this answer from [Helmenstine2019a].

The association constant, $K_a$,

$$Ka = [H+][B-]/[HB] \quad (S1)$$

where $[H^+]$ is concentration of $H^+$ ions, $[B^-]$ is concentration of conjugate base ions, and [HB] is the concentration of undissociated acid molecules. We assume that the acid releases one $H^+$ ion for every $B^-$ ion, so

$$[H^+] = [B^-]. \quad (S2)$$

To simplify the algebra, we denote $[H^+]$ as *x.* Thus,

$$[HB] = C - x,$$

where *C* is the analytical concentration of the acid. Thus, using **Equation S1**,

$$K_a = \frac{x \times x}{C - x}, \quad (S3)$$

$$x^2 = K_a (C - x), \quad (S4)$$

$$x^2 + K_a x - C K_a = 0. \quad (S5)$$

The proton concentration, *x,* can be found using the positive root of the quadratic equation,

$$x = -\frac{b}{2a} \pm \frac{1}{2a}\sqrt{b^2 - 4ac}, \quad (S6)$$

$$x = -\frac{K_a}{2} \pm \frac{1}{2}\sqrt{K_a^2 + 4C K_a}. \quad (S7)$$

Thus,

$$pH = -\log_{10}\left(-\frac{K_a}{2} + \frac{1}{2}\sqrt{K_a^2 + 4C K_a}\right). \quad (S8)$$

For acetic acid (pKa = 4.75), citric acid (pKa = 3.13), and gluconic acid (pKa = 3.72) at 100 mM,

$$pH_{acetic} = 2.9, \quad (S9)$$

$$pH_{gluconic} = 2.4, \quad (S10)$$

$$pH_{citric} = 2.1. \quad (S11)$$

### Supplementary Information References

[Alissandratos2015a] A. Alissandratos and C. J. Easton. "Biocatalysis for the application of $CO_2$ as a chemical feedstock". *Beilstein Journal of Organic Chemistry* 11 (2015), pp. 2370–2387. doi:10.3762/bjoc.11.259.

[Bar-Even2016a] A. Bar-Even. "Formate Assimilation: The Metabolic Architecture of Natural and Synthetic Pathways." *Biochemistry* 55 (2016), pp. 3851–63. doi:10.1021/acs.biochem.6b00495.

[Barstow2021b] B. Barstow. "ElectroCO$_2$". *Zenodo* (2021). doi:10.5281/zenodo.5805345.

[Berg2002a] J. Berg, J. Tymoczko, and L. Stryer. Biochemistry. 5th. New York, NY: W H Freeman, 2002.

[Berg2011a] I. A. Berg. "Ecological Aspects of the Distribution of Different Autotrophic $CO_2$ Fixation Pathways". *Applied and Environmental Microbiology* 77 (2011), pp. 1925–1936. doi:10.1128/aem.02473-10.

[BergI2007a] I. A. Berg, D. Kockelkorn, W. Buckel, and G. Fuchs. "A 3-Hydroxypropionate/4-Hydroxybutyrate Autotrophic Carbon Dioxide Assimilation Pathway in Archaea". *Science* 318 (2007), pp. 1782–1786. doi:10.1126/science.1149976.

[Claassens2016a] N. J. Claassens, D. Z. Sousa, V. A. M. dos Santos, W. M. de Vos, and J. van der Oost. "Harnessing the power of microbial autotrophy". *Nature Reviews Microbiology* 14 (2016), pp. 692–706. doi:10.1038/nrmicro.2016.130.

[Helmenstine2019a] T. Helmenstine, "How to Calculate the pH of a Weak Acid". https://www.thoughtco.com/calculating-ph-of-a-weak-acid-problem-609589 (Visited on 01/08/2022).

[Herter2002a] S. Herter, G. Fuchs, A. Bacher, and W. Eisenreich. "A bicyclic autotrophic $CO_2$ fixation pathway in *Chloroflexus aurantiacus*". *Journal of Biological Chemistry* 277 (2002), pp. 20277–20283. doi:10.1074/jbc.m201030200.

[Huber2008a] H. Huber, M. Gallenberger, U. Jahn, E. Eylert, I. A. Berg, D. Kockelkorn, W. Eisenreich, and G. Fuchs. "A dicarboxylate/4-hydroxybutyrate autotrophic carbon assimilation cycle in the hyperthermophilic *Archaeum Ignicoccus hospitalis*". *Proceedings of the National Academy of Sciences* 105 (2008), pp. 7851–7856. doi:10.1073/pnas.0801043105.

[Kanehisa2000a] M. Kanehisa and S. Goto. "KEGG: Kyoto Encyclopedia of Genes and Genomes". *Nucleic Acids Research* 28.1 (2000), pp. 27–30. doi:10.1093/nar/28.1.27.

[Kanehisa2019a] M. Kanehisa. "Toward understanding the origin and evolution of cellular organisms". Protein *Science* 28.11 (2019), pp. 1947–1951. doi:10.1002/pro.3715.

[Kanehisa2021a] M. Kanehisa, M. Furumichi, Y. Sato, M. Ishiguro-Watanabe,and M. Tanabe. "KEGG: integrating viruses and cellular organisms". *Nucleic Acids Research* 49.D1 (2020), gkaa970–. doi:10.1093/nar/gkaa970.

[Salimijazi2020b] F. Salimijazi, J. Kim, A. M. Schmitz, R. Grenville, A. Bocarsly, and B. Barstow. "Constraints on the Efficiency of Engineered Electromicrobial Production". *Joule* 4 (2020), pp. 2101–2130. doi:10.1016/j.joule.2020.08.010.

[Schomburg2017a] I. Schomburg, L. Jeske, M. Ulbrich, S. Placzek, A. Chang,and D. Schomburg. "The BRENDA enzyme information system–From a database to an expert system". *Journal of Biotechnology* 261 (2017), pp. 194–206. doi:10.1016/j.jbiotec.2017.04.020.

[Siegel2015a] J. B. Siegel, A. L. Smith, S. Poust, A. J. Wargacki, A.Bar-Even, C. Louw, B. W. Shen, C. B. Eiben, H. M.Tran, E. Noor, J. L. Gallaher, J. Bale, Y. Yoshikuni, M. H. Gelb, J. D. Keasling, B. L. Stoddard, M. E. Lidstrom, and D. Baker. "Computational protein design enables a novel one-carbon assimilation pathway". *Proceedings of the National Academy of Sciences* 112 (2015), p. 3704-3709. doi:10.1073/pnas.1500545112.

[Zarzycki2009a] J. Zarzycki, V. Brecht, M. Müller, and G. Fuchs. “Identifying the missing steps of the autotrophic 3-hydroxypropionate $CO_2$ fixation cycle in *Chloroflexus aurantiacus*”. *Proceedings of the National Academy of Sciences* 106 (2009), p. 21317. doi.10.1073/pnas.0908356106.
